# All-or-none visual categorization in the human brain

**DOI:** 10.1101/658625

**Authors:** Talia L. Retter, Fang Jiang, Michael A. Webster, Bruno Rossion

## Abstract

Whether visual categorization, i.e., specific responses to a certain class of visual events across a wide range of exemplars, is graded or all-or-none in the human brain is largely unknown. We address this issue with an original frequency-sweep paradigm probing the evolution of responses between the minimum and optimal presentation times required to elicit both neural and behavioral face categorization responses. In a first experiment, widely variable natural images of nonface objects are progressively swept from 120 to 3 Hz (8.33 to 333 ms duration) in rapid serial visual presentation sequences; variable face exemplars appear every 1 s, enabling an implicit frequency-tagged face-categorization electroencephalographic (EEG) response at 1 Hz. In a second experiment, faces appear non-periodically throughout such sequences at fixed presentation rates, while participants explicitly categorize faces. Face-categorization activity emerges with stimulus durations as brief as 17 ms for both neural and behavioral measures (17 – 83 ms across individual participants neurally; 33 ms at the group level). The face-categorization response amplitude increases until 83 ms stimulus duration (12 Hz), implying graded categorization responses. However, a strong correlation with behavioral accuracy suggests instead that dilution from missed categorizations, rather than a decreased response to each face stimulus, may be responsible. This is supported in the second experiment by the absence of neural responses to behaviorally uncategorized faces, and equivalent amplitudes of isolated neural responses to only behaviorally categorized faces across presentation rates, consistent with the otherwise stable spatio-temporal signatures of face-categorization responses in both experiments. Overall, these observations provide original evidence that visual categorization of faces, while being widely variable across human observers, occurs in an all-or-none fashion in the human brain.

## Introduction

A fundamental function of nervous systems is to organize the flurry of sensory inputs generated by their rich, dynamic, and ambiguous environment. This organization requires generating distinct responses to different sensory inputs (i.e., discrimination), but also generating a common response to different but related sensory inputs (i.e., generalization). This categorization function plays a fundamental role in responding adaptively to the environment, and has been considered to be the foundation of many other cognitive functions, allowing us to learn, memorize, act, and communicate through language, gesture, and expression (Smith & Medin, 1981; Edelman, 1987; Murphy, 2002; Goldstone, Kersten & Carvalho, 2018).

An outstanding issue is whether neural categorization of currently experienced stimuli (i.e., perceptual categorization) is graded or all-or-none. That is, does our brain progressively build categorization responses, or does categorization emerge all at once from an accumulation of non-categorical sensory information? This issue is important insofar as perceptual categorization relates to our subjective experience of the stimulus, or perceptual awareness (e.g., del Cul et al., 2007; Quiroga et al., 2008; de Gardelle et al., 2011; Bachmann, 2013; Sekar et al., 2013; Navajas et al., 2014; Windey et al., 2015). Understanding whether neural categorization is all-or-none may constrain psychological, philosophical, and computational constructs of conscious awareness, as well as advance theoretical models of brain function.

Studies that have addressed this issue have generally relied on vision, the dominant modality in humans and other primates, and used neural measures recorded during variable stimulus viewing conditions. On one hand, a number of studies have found that neural activity in visual cortical areas increases in an analog, continuous way with increases in stimulus visibility (Kovács, Vogels & Orban, 1995; Vanni et al, 1996; Grill-Spector et al, 2000; Keysers et al, 2001; Bar et al., 2001; Moutoussis & Zeki, 2002; Bacon-Macé et al, 2005; Christensen et al., 2006), supporting a graded or progressive visual categorization process. On the other hand, other studies have attempted to relate neural activity to explicit behavioral reports and rather concluded in favor of all-or-none brain responses related to perceptual awareness, more like discrete “hits” and “misses” (Del Cul et al., 2007; Quiroga et al., 2008; Harris, Wu & Woldorff, 2011; Shafto & Pitts, 2015). The abundance of evidence on both sides has been difficult to reconcile, largely hindering rather than inspiring a coherent theoretical framework.

To address this fundamental yet unresolved issue in cognitive neuroscience (Windey & Cleeremans, 2015), we introduce an approach that goes beyond previous efforts to address this question in a number of important ways. First, going beyond previous studies that compared to detection or identification of a limited set of specific stimuli without exemplar generalization, we truly measure neural *categorization*, i.e., responses that are specific to a certain class of stimuli across a wide range of variable exemplars. To do that, we use a large set of widely variable natural images of faces. Faces are used as the visual stimuli to categorize for a number of reasons. From the first minutes of life, the face is arguably the most important visual stimulus for human ecology. Faces are complex multidimensional stimuli that are ubiquitous in our visual environment and drive many of our behaviors. Their initial categorization as “a face” (i.e., generic face categorization) unfolds into an extremely rich social categorization of the individual, allowing to categorize people according to their gender, ethnicity, emotional expression, age, attractiveness, and identity. In neurotypical human adults, generic face categorization evokes extensive neural activity along the (ventral) occipital temporal cortex (e.g., Sergent, Ohta & MacDonald, 1992; Allison et al., 1994; Puce et al., 1995; Kanwisher, McDermott & Chun, 1997; Weiner & Grill-Spector, 2010; Zhen et al., 2015; Jonas et al., 2016; Jacques et al., 2016; Gao et al., 2018). As shown with face Pareidolia, generic face categorization in humans goes well beyond the identification of well-defined objective physical features of real faces (Caharel et al., 2013; Omer et al., 2019), providing an advantage over artificial systems in terms of generalization within natural and degraded views and environments (Scheirer et al., 2014).

Second, we relate human’s (implicit) neural and (explicit) behavioral responses always under the same stimulation constraints, each face exemplar being immediately preceded and followed by a nonface object, i.e. visually masked (Helmholtz, 1867; Enns & Di Lollo, 2000). Compared to studies that have used a backward-masking approach to probe perceptual awareness (e.g., Del Cul et al., 2007; Quiroga et al., 2008; Sekar, Findley & Linas, 2012; Sekar et al., 2013), we use both backward- and forward-masking. Most importantly, our stimuli are presented in a dynamic visual stream, i.e., variable faces embedded in a rapid train of variable nonface object images for about 1 minute (Figure 1), similarly to rapid serial visual presentation (RSVP) sequences (Potter & Levy, 1969; Keysers et al., 2001; 2005; Potter et al., 2014). This paradigm, combined with neural frequency-tagging (Adrian & Matthews, 1934; Regan, 1989; Norcia et al., 2015) is known to simultaneously measure the two key aspects of generic face categorization, namely *discrimination* by measuring selective responses to face stimuli embedded in rapid trains of non-face objects, and *generalization* by measuring the common selective responses to widely variable exemplars of human faces (Rossion et al., 2015). The neural face categorization responses obtained are devoid of low-level confounds (i.e., amplitude spectrum; Rossion et al., 2015; Gao et al., 2018), and associated with robust neural activity in face-selective regions of the ventral occipito-temporal cortex as shown both with human intracerebral recordings (Jonas et al., 2016) and functional magnetic resonance imaging (Gao et al., 2018).

**Figure 1.**
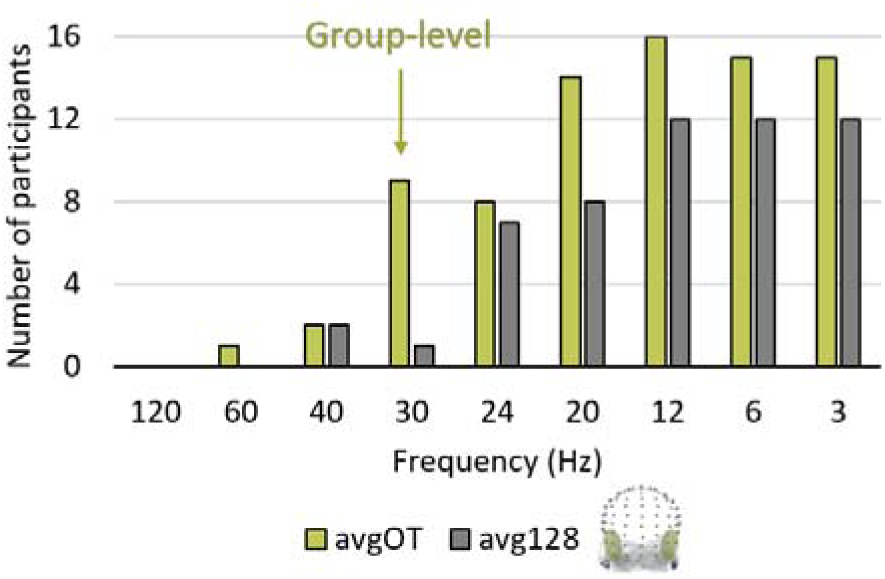
The number of participants (out of 16 in total) with significant responses at each frequency rate (Z > 1.64; p< .05; Experiment 1). At the group-level, the response was first significant at 30 Hz over the occipito-temporal ROI (Z = 3.5, p = .0002). **Key)** avgOT = average across the left and right occipito-temporal ROIs; avg128 = average of all 128 EEG channels.

Here, to probe the all-or-none or graded nature of face categorization, we increasingly sweep stimulus duration in this paradigm, from 120 Hz stimulation rate (i.e., 8.33 ms SOA) to 3 Hz (333 ms SOA). We then track the onset, optimum and interindividual variability of the neural face categorization response. Importantly, in this case categorization is measured implicitly, i.e., without requiring explicit categorization judgments from the observers, another significant difference with previous studies. However, we also collect explicit behavioral measures at different times under the same stimulation conditions, and relate the two measures on an individual basis.

As we document below, our observations indicate that while general visual responses can be tagged at presentation rates up to 120 Hz over the medial occipital cortex (Herrmann, 2001), face categorization emerges significantly as early as 60 Hz (17 ms SOA/duration) for a few trials, then reliably at the group level at 30 Hz (i.e., 33 ms duration in a train of nonface objects) to saturate at 12 Hz (83 ms duration) over occipito-temporal regions. However, there is substantial interindividual variability in the presentation rates for face categorization responses (emerging between 17 and 83 ms for different observers), and importantly, these differences are correlated across behavioral and neural measures. Crucially, our findings reveal that the apparent graded increase in the neural amplitudes of face-categorization responses in fact reflects an all-or-none response occurring occasionally to faces at extremely short viewing times and becoming more consistent, but not more evolved, as presentation duration increases. The tight relationship with explicit behavior indicates that this all-or-none face categorization response corresponds to perceptual awareness.

## Materials and Methods

### Participants

Sixteen adult participants at the University of Nevada, Reno took part in the experiment All reported normal or corrected-to-normal visual acuity. Their mean age was 29 years (SE = 1.65; range = 22 to 44), ten identified as female and six as male, and 15 were right-handed. The study was conducted in accordance with the Code of Ethics of the World Medical Association (Declaration of Helsinki), with protocols approved by the University of Nevada Institutional Review Board.

### Stimuli and Display

Stimuli were 112 natural face images and 200 natural object images compiled internally (for examples, see Movie 1; Quek et al., 2018; Gao et al., 2018). Facial identities were selected to represent a broad range of characteristics (e.g., age, sex, expression, head orientation). Both face and object images were edited from photographs taken under variable environmental conditions and perspectives (in lighting, background, orientation, etc.). Each image was cropped to a square with the target face or object centered or off-center, resized to 200×200 pixels, and saved in JPEG format. Stimulus variations were not further controlled because the high amount of variability within both face and object categories reduces the diagnosticity of these cues to any one category (Thorpe et al., 1996; Crouzet et al., 2010; Foldiak et al., 2004; Rossion et al., 2015). The stimuli were presented on a cathode ray tube (CRT) monitor (NEC AccuSync 120) with a 120 Hz screen refresh rate and a resolution of 800×600 pixels. At the viewing distance of 80 cm, stimuli subtended 7.3° by 7.3° degrees of visual angle.

### Experiment 1 (EEG Frequency-Tagging) Procedure

After the setup of the EEG system, participants were positioned in front of the monitor, with access to a keyboard with which to start stimulation trials and give responses. The only illumination in the room came from the testing and acquisition monitors. Participants were informed that they would see series of natural images presented at a high frequency rate, which would reduce incrementally throughout each trial. They were not given any information about the types of images they would see. Instead, they were instructed to fixate on a centrally presented fixation cross throughout the experiment, and to press on the space bar when the cross briefly changed luminance, from black to white, 12 times per trial.

Each trial consisted of: 1) the cross to establish fixation lasting between 2-4 s; 2) a 63- s sequence, consisting of nine, contiguous 7-s steps of stimulus presentation at 120, 60, 40, 30, 24, 20, 12, 6, and then 3 Hz; and finally, the fixation cross for another 2-4 s. Images were presented continuously with a 100% duty cycle at maximum contrast (Retter et al., 2018). Frequency rates followed integer multiples of the 120 Hz monitor frame rate, i.e. with 1 frame at 120 Hz (8.3 ms), 2 frames at 60 Hz (17 ms), 3 frames at 40 Hz (25 ms), etc. Throughout each trial, image size varied randomly at each presentation between 95-105% of the original in five steps.

Crucially, regardless of the frequency of image presentation throughout the sequence, faces were embedded consistently at a rate of 1 Hz. Thus, for example, at 120 Hz, faces appeared as every one out of 120 images, while at 6 Hz faces appeared as every one out of 6 images. The ratio of face to object images has been shown not to have an impact on the face-categorization response, when as here the faces are separated by at least 400 ms (Retter & Rossion, 2016a). The order of face stimuli and object stimuli was randomized for each subject and trial repetition. Participants binocularly viewed 18 trial repetitions, leading to a total testing time of about 30 minutes.

### Experiment 2 (Behavior and ERP) Procedure

After Experiment 1, participants were asked two questions: 1) “What did you notice about the images you just saw?”; and 2) “Did you notice any periodicity within the sequences?” In response to the first, open-ended question, participants typically reported seeing a lot of images, the rate of presentation decreasing, and that they noticed faces (14 participants), animals (10 participants), non-living objects (8 participants), and plants/natural scenes (5 participants). Some participants reported one or a few specific images reoccurring: e.g., a dog, elephant, clock, elderly woman, or car. Importantly, in response to the second question, no participants reported noticing any periodicity of faces within the sequences.

Participants were then debriefed in regards to the first experiment, including being informed that faces had appeared every 1 s throughout all of the sequences. They were then given instructions for the second experiment, which instead involved an explicit face detection task. Participants were asked to respond to human faces, which would appear non-periodically, or not at all, within shorter (30 s) sequences, each at a single frequency throughout, by pressing on the key “J” with the index finger of their dominant hand. The faces appeared 5-7 times in five sequences per condition, and 0 times in one sequence per condition. This resulted in a total of about 30 face presentations for each of the nine frequency conditions (M = 30.4; SD = 2.89), for a total duration of about 40 minutes.

### EEG Acquisition

EEG acquisition and analysis modeled protocols established in previous studies (e.g., Jacques, Retter & Rossion, 2016; Retter & Rossion, 2016a; Retter et al., 2018). A BioSemi ActiveTwo system (BioSemi B.V., Amsterdam, The Netherlands) with 128 Ag-AgCl Active-electrodes was used to record the EEG. Electrodes were organized in the default BioSemi configuration, centered around nine standard 10/20 locations on the cardinal axes (for exact position coordinates, see http://www.biosemi.com/headcap.htm). For ease of comparison across studies, the default electrode labels were updated to closely match the conventional 10/5 system (for exact relabeling, see Rossion et al., 2015, Figure S2; Oostenveld & Praamstra, 2001). Additionally, vertical and horizontal electrooculogram (EOG) were recorded with four flat-type Active-electrodes: two above and below the right eye and two lateral to the external canthi. The offset of all electrodes was set below 50 mV, relative to the additional common mode sense (CMS) and driven right leg (DRL) electrode loop. The EEG and EOG were digitized at a sampling rate of 2048 Hz and subsequently downsampled offline to 512 Hz.

### Analysis

Letswave 5, an open source toolbox (http://nocions.webnode.com/letswave) running over MATLAB R2013b (The MathWorks, USA), was used for the data analysis.

#### Preprocessing

Data were bandpass filtered from 0.05 to 140 Hz with a fourth-order zero-phase Butterworth filter. Sequences were coarsely segmented from 2 s before to 65 s after sequence onset. To correct for blinks, an independent component analysis was applied for the six participants who blinked more than 0.2 times/s on average throughout the testing sequences (M = 1.20 blinks/s; SD = 0.22; as in, e.g., Retter & Rossion, 2016a). Channels with abrupt deflections of ± 100 µV or above in more than one epoch were linearly interpolated with the neighboring 3-4 channels (M = 0.75 channels; SE = 0.86 channels; range = 0-3 channels, i.e., up to no more than 3% of channels).

Data were re-referenced to the average of all 128 EEG channels. Each 7-s presentation duration step was re-segmented in two ways: 1) beginning from −200 ms before the first face stimulus onset, to capture face-categorization responses; and; 2) beginning exactly at face onset, in order to isolate the stimulus-presentation response to the full 7 s of each presentation duration step.

#### Regions-of-interest

To determine significance of the face-categorization responses, a ten-channel bilateral occipito-temporal (OT) ROI was defined a priori based on previous studies with the generic face categorization paradigm (e.g., Retter & Rossion, 2016b; Retter et al., 2018; see also Dzhelyova & Rossion, 2014). To decompose the responses further, the average amplitude of the right and left OT regions were examined separately (right: channels P10; P8; PO8; PO10; PO12; and left: P9; P7; PO7; PO9; PO11). The bilateral ROI was verified to capture face-categorization responses post-hoc (see Results). To provide a region-free assessment of response significance, we also investigated responses across the average of all 128 EEG channels. Finally, to identify and quantify face-categorization responses, we used a medial-occipital (MO) region consisting of the 10 following channels: O2; POI2; I2; Iz; OIz; Oz; POOz; O1; POI1; I1), as well as the 128-channel grand average.

#### Frequency Domain Analysis

The segmented data were averaged by presentation duration step in the time domain, reducing non-phase locked (i.e., non-stimulus related) activity and thereby increasing the signal-to-noise ratio. A fast Fourier transform (FFT) was applied to represent the data of each channel in the temporal frequency domain, i.e., as a normalized amplitude spectrum (µV), with a range of 0 to 256 Hz and a resolution of 0.14 Hz. To correct for variations in baseline noise level around each frequency of interest (i.e., 1 Hz and its harmonics for face-categorization responses, and the stimulus presentation rate and its harmonics), the average amplitude of the neighboring six frequency bins were averaged (a range of 0.86 Hz) and subtracted from each frequency bin. The harmonics of the face-categorization responses were subsequently summed from the fundamental, 1 Hz, up to 20 Hz (similarly to Retter & Rossion, 2016a; Retter et al., 2018), excluding harmonics that coincided with the stimulus presentation frequency when present. The harmonics of stimulus-presentation responses were summed from the fundamental up to 120 Hz for all presentation duration steps.

To assess significance of the face-categorization and stimulus-presentation responses at each presentation duration condition at the group level, a Z-Score was computed on the average of the bilateral OT ROI channels for face-categorization responses, and the average of the MO ROI channels for the stimulus-presentation response (using the same baseline frequency range as for the baseline-subtraction). Additionally, the same approach was used at the individual participant level. Responses were considered to be significant when the Z- Score exceeded 2.32 (p<0.01, 1-tailed, i.e., predicting the signal was greater than the noise).

To describe the data further, the amplitude and scalp topographies were assessed across presentation conditions. Scalp topographies were visualized in terms of both their original amplitude and their normalized amplitude (McCarthy & Wood, 1985). To assess the extent of right lateralization for face categorization, a lateralization index was computed using the amplitudes of the right and left OT ROIs (R and L, respectively) as follows: (R-L)/(R+L)*100 (as in Retter & Rossion, 2016a). Statistical tests on response amplitudes were applied independently for face-categorization and stimulus-presentation responses in the form of repeated measures analysis-of-variance tests (ANOVAs), with factors of *Region* and *Condition*. A Greenhouse-Geisser correction was applied to the degrees of freedom in the event that Mauchly’s test of sphericity was significant. Data were grand-averaged across participants for display purposes.

#### Time Domain Analysis

Segmented data were filtered more conservatively with a fourth-order, zero-phase Butterworth low-pass filter at 30 Hz. A notch filter was applied to remove the periodic responses to stimulus-presentation for each presentation condition at its fundamental and harmonic frequencies up to 30 Hz. Data were then re-segmented as described previously for face-categorization responses and averaged by condition. A baseline correction of voltage offset was applied on the 200 ms immediately preceding face stimulus onset.

To test for statistical significance of the response from the baseline reference interval (−200 to 0 ms) at each time point, bootstrapping was performed with a significance criterion of p<0.001, with 10,000 iterations. To reduce the likelihood of false positives, only groups of at least 5 consecutive time bins (i.e., about 10 ms) were reported. Grand-averaging was computed for display. Finally, to describe a face-categorization response independent of the stimulus presentation duration, the data were averaged across all presentation-duration conditions yielding significant frequency-domain face-categorization responses.

#### Behavioral Face-Categorization Analysis (Experiment 2)

Face-categorization responses were considered correct when occurring between 0.15 – 2 s after face presentation onset; responses occurring outside this range were considered false positives. This criterion rejected 3.5% of potentially correct face-categorization responses, and generated a response time distribution with a similar mean and median: 509 (SE = 117.5 ms) and 512 ms, respectively. False positives occurred on average at a rate of 0.24 per minute (SE = 0.087); they occurred maximally in a mid-frequency range, from 20–40 Hz, although this may be for several reasons (e.g., that participants found the task too difficult to try at the highest frequencies, or that they actually did perceive faces sometimes in the stream of non-face objects (pareidolia); Supplemental Figure 3). Note that this generous window for correct responses was chosen in terms of the minimum and maximal expected response times for any participant for any key press (the minimal time between two consecutive face images was 2.6 s); however, application of a more conservative range from 0.2–1 s after face presentation only minimally affected the pattern of results across participants or frequencies (e.g., Supplemental Figure 3, for the effect on false positives).

The total percent accuracy was defined as the percent of correct detection responses minus the percent of false positives, relative to the total number of faces presented. This modification of the raw accuracy guarded against the chance that a high rate of correct responses occurred as a byproduct of a high rate of false positives; this was particularly important given the differences in false alarms across presentation rates. Response time (RT) was considered only for correct responses.

Additionally, inverse efficiency (IE) was computed by dividing RT by total percent accuracy for each participant (Townsend & Ashby, 1983). This IE measure, combining accuracy and RT, was used to rank participants in their face-categorization ability overall, by using the average of trials from all presentation conditions together. Note that IE is thus used only to rank participants’ face-categorization *ability*, while in all other analyses accuracy is used in order to examine *whether or not* faces were categorized, irrespective of the RT. The relationship between behavioral face-detection ability and EEG responses was tested with Pearson correlations between IE and EEG response amplitude. Differences in correlation coefficient strength were calculated according to the asymptotic z-test of Lee and Preacher (2013).

#### Behavioral Fixation-Detection Analysis (Experiment 1)

Responses to the detection of the fixation cross luminance change were analyzed with the same parameters as for the face-categorization responses. Overall, participants’ accuracy was high for this easy task across all frequencies (93.4%; SE = 1.39%), with a mean correct RT of 480.0 ms (SE = 16.27 ms). The rate of false positives was low: 0.15 per minute (SE = 0.038). The accuracy was higher at the highest frequencies (above 96% at 30 Hz and above; relative to 91–93% at all lower frequencies), although this could be either because the luminance changes of the cross were easier to see against a more rapid (i.e., less well-processed) image stream, or as an order effect, with the higher frequencies consistently appearing first in the sequences. Overall, the fixation cross task results differed greatly from those of the face categorization task, emphasizing the differences in these tasks and processes.

## Results

The emergence of face-categorization responses (Section 1) was first determined at a neural level from the EEG responses to periodic faces in Experiment 1 in the frequency domain, and at a behavioral level from the face-categorization response accuracy to non-periodic faces in Experiment 2. Then, these neural and behavioral face-categorization responses were quantified across presentation rates from 120 to 3 Hz (Section 2). In Section 3, the relationship between neural and behavioral face-categorization responses is characterized across presentation rates, and explained in terms of neural responses (event-related potentials) to behaviorally categorized vs. uncategorized faces in Experiment 2. A qualitative assessment of face-categorization responses across presentation rates is presented in Section 4. Finally, individual differences in generic face categorization are investigated in Section 5.

### 1. Emergence of Face-Categorization Responses

#### 1.1 Neural Responses (Experiment 1)

At the individual participant level, significant summed-harmonic OT face-categorization responses emerged between a range of 60 to 23 Hz (17 to 83 ms; Figure 1). On the high end, a single participant had a significant response at 60 Hz, and two participants followed at 40 Hz. On the low end, a single participant had a significant response only up to 12 Hz, while three participants had responses up to 20 Hz. The mean across participants was 29.4 Hz (SE 12 = 2.73 Hz), and the median 30.0 Hz.

On average across observers, face-categorization responses appeared to first emerge at 40 Hz (25 ms) over occipito-temporal channels (Figure 2A), although the response was not significant until 30 Hz (33 ms) over the OT ROI (Table 1A; see also Table 2A). When considering response significance from the average of all 128 EEG channels, the highest significant face-categorization response was reduced to 20 Hz (50 ms). In contrast, the stimulus-presentation responses were significant at all frequencies tested at the group level, both at the MO ROI and the average of all 128 channels (Table 1B).

**Table 1.** Group-level Z-scores, calculated at the occipito-temporal (OT) ROI for face-categorization responses (**A**), the medial-occipital (MO) ROI for stimulus-presentation responses (**B**), and the average of all 128 EEG channels for both face- and stimulus-presentation responses. Significant responses are shown in bold (Z > 2.32, p<.01).

| Frequency (Hz) | 120 | 60 | 40 | 30 | 24 | 20 | 12 | 6 | 3 |
| --- | --- | --- | --- | --- | --- | --- | --- | --- | --- |
| **A. Face** |  |  |  |  |  |  |  |  |  |
| OT | 0.11 | -0.14 | 1.12 | **3.50** | **5.09** | **6.77** | **10.37** | **8.87** | **6.92** |
| Avg128 | 0.25 | 0.04 | 0.25 | 0.55 | 1.46 | **2.37** | **3.02** | **3.23** | **2.38** |
| **B. Stimulus** |  |  |  |  |  |  |  |  |  |
| MO | **14.73** | **24.33** | **49.19** | **72.08** | **50.22** | **98.41** | **71.87** | **328.60** | **63.68** |
| Avg128 | **18.16** | **18.68** | **22.51** | **53.25** | **30.01** | **74.36** | **47.62** | **144.46** | **35.99** |

**Table 2.** Mean response amplitude (µV), calculated at the occipito-temporal (OT) and medial-occipital (MO) ROIs for both face-categorization (**A**) and stimulus-presentation (**B**). One standard error, across participants, is included in parentheses.

| Frequency (Hz) | 120 | 60 | 40 | 30 | 24 | 20 | 12 | 6 | 3 |
| --- | --- | --- | --- | --- | --- | --- | --- | --- | --- |
| **A. Face ($\mu V$)** |  |  |  |  |  |  |  |  |  |
| OT | 0.03<br>(0.06) | -0.03<br>(0.08) | 0.21<br>(0.08) | 0.57<br>(0.12) | 0.93<br>(0.23) | 1.43<br>(0.21) | 1.95<br>(0.25) | 1.66<br>(0.21) | 1.72<br>(0.19) |
| MO | 0.008<br>(0.055) | 0.07<br>(0.07) | 0.22<br>(0.09) | 0.26<br>(0.08) | 0.47<br>(0.09) | 0.78<br>(0.13) | 1.06<br>(0.18) | 1.05<br>(0.12) | 1.10<br>(0.18) |
| **B. Stimulus ($\mu V$)** |  |  |  |  |  |  |  |  |  |
| OT | 0.04<br>(0.005) | 0.05<br>(0.006) | 0.13<br>(0.01) | 0.22<br>(0.02) | 0.24<br>(0.03) | 0.35<br>(0.05) | 0.59<br>(0.06) | 1.62<br>(0.13) | 2.40<br>(0.18) |
| MO | 0.03<br>(0.004) | 0.06<br>(0.008) | 0.16<br>(0.01) | 0.33<br>(0.03) | 0.45<br>(0.04) | 0.64<br>(0.08) | 1.11<br>(0.12) | 2.05<br>(0.19) | 3.35<br>(0.27) |

#### 1.2 Behavioral Responses (Experiment 2)

At the group level (Figure 2B; Table 3A), behavioral face-categorization accuracy, corrected for false-positives, emerged at 60 Hz (17 ms; M = 3.03%; SE = 1.26%), t_15_ = 2.41, p = .015, one-tailed, d = 0.85. At the higher frequency step, 120 Hz, accuracy did not differ from 0% (−0.40%; SE = 0.50%); at the subsequent frequency step, 40 Hz, accuracy jumped to 22% (SE = 3.2%).

**Figure 2.**
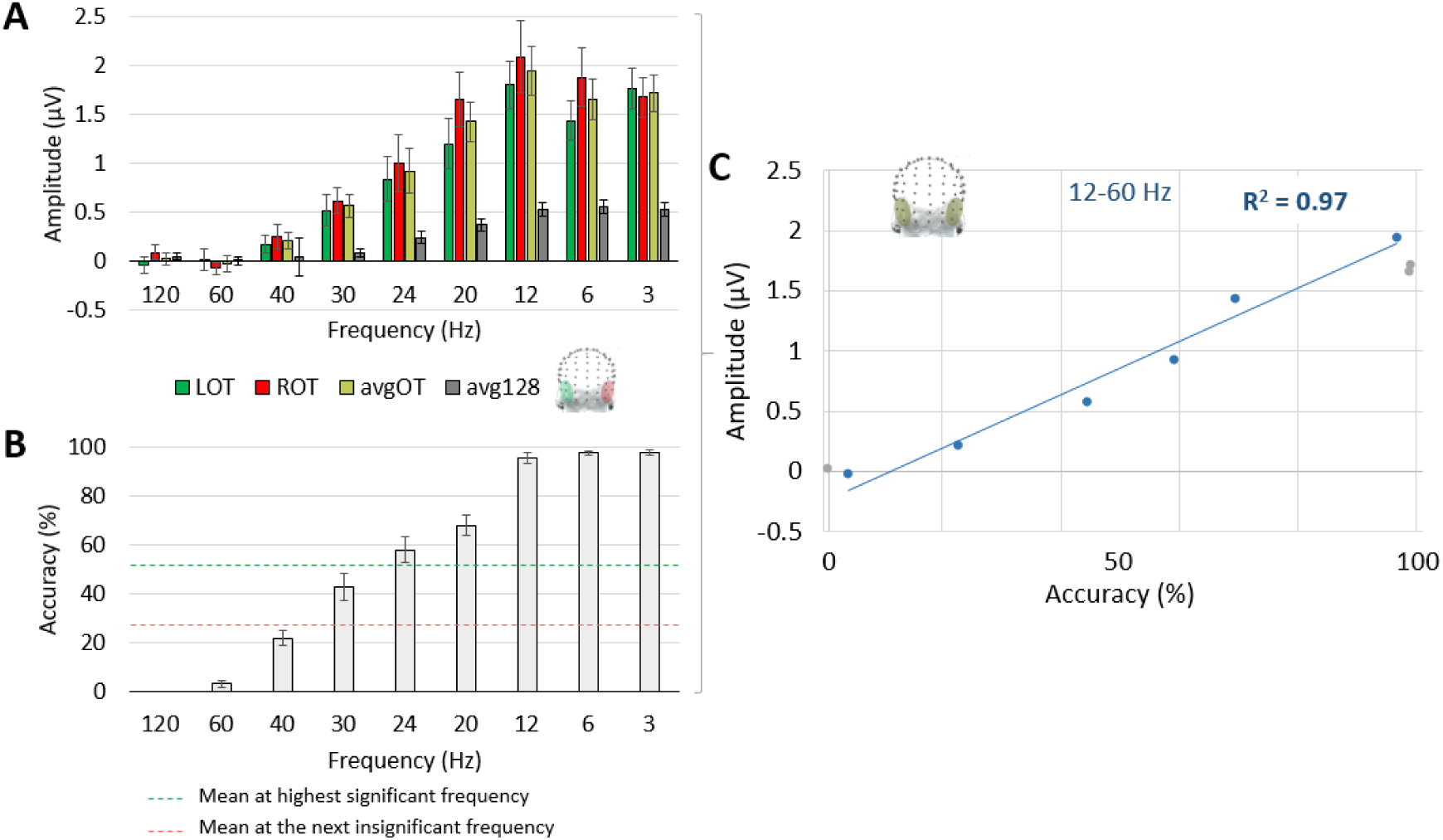
Behavioral face-categorization accuracy (Experiment 2) compared with neural face-categorization response amplitude (Experiment 1). **A**) Baseline-subtracted summed-harmonic amplitudes across different ROIs. **B**) Behavioral mean face-categorization accuracy across frequency rates. The dashed green line at 52% accuracy represents the mean accuracy corresponding to highest frequency yielding a significant face categorization response for each participant; the dashed red line at 27% represents the mean accuracy corresponding to the frequency one step higher for each participant, at which the face categorization response was not significant. **C)** A linear relationship between accuracy (Experiment 2) and amplitude (Experiment 1) between 12 and 60 Hz. The data points from the remaining presentation rates (3, 6, and 120 Hz) are shown in light gray, but were not included in this correlation. **Key)** LOT: left occipito-temporal ROI; ROT: right occipito-temporal ROI; avgOT = average of these left and right occipito-temporal ROIs; avg128 = average of all 128 EEG channels.

**Table 3.** Behavioral face-detection results: percent accuracy (**A**), response time (RT; **B**), and inverse efficiency (IE; **C**). One standard error for each measurement is given in parentheses.

| Frequency (Hz) | 120 | 60 | 40 | 30 | 24 | 20 | 12 | 6 | 3 |
| --- | --- | --- | --- | --- | --- | --- | --- | --- | --- |
| **A. Accuracy (%)** | -0.4<br>(0.49) | 3.0<br>(1.26) | 21.7<br>(3.17) | 42.9<br>(5.59) | 57.9<br>(5.20) | 68.0<br>(4.32) | 95.6<br>(2.12) | 97.8<br>(0.86) | 98.0<br>(1.01) |
| **B. RT (ms)** | 1435<br>(145) | 650<br>(52.7) | 557<br>(19.2) | 529<br>(18.9) | 505<br>(11.0) | 503<br>(15.2) | 498<br>(13.4) | 495<br>(15.3) | 499<br>(15.2) |
| **C. IE** | 84.7<br>(260) | 74.5<br>(20.8) | 32.0<br>(4.85) | 14.2<br>(1.91) | 10.1<br>(1.16) | 8.11<br>(0.81) | 4.29<br>(0.26) | 5.07<br>(0.18) | 5.11<br>(0.18) |

### 2. Quantization of Face-Categorization Responses Across Presentation Rates

#### 2.1 Neural Responses (Experiment 1)

The amplitude of face-categorization responses was quantified at four ROIs: over the OT ROI, as well as its divisions into the right occipito-temporal (ROT) and left occipito-temporal (LOT) ROIs, and the average of all 128 channels (Figure 2A). Across quantification regions, the response amplitude neared zero at 120 and 60 Hz, and then increased progressively until 12 Hz (83 ms; Figure 3B). After 12 Hz, it decreased but remained significant until the slowest presentation rate, 3 Hz (333 ms; Table 1A). In comparison, the amplitude of the stimulus-presentation response decreased continuously from 3 to 120 Hz, but always remained above zero (Supplemental Figure 1). The face-categorization response was at least about 3 times larger over the OT than average 128- channel region over all conditions below 60 Hz, indicating a relatively focal response (for stimulus presentation, the MO ROI captured a lesser proportion of the response, on average a little over 2 times larger than the 128-channel average).

**Figure 3.**
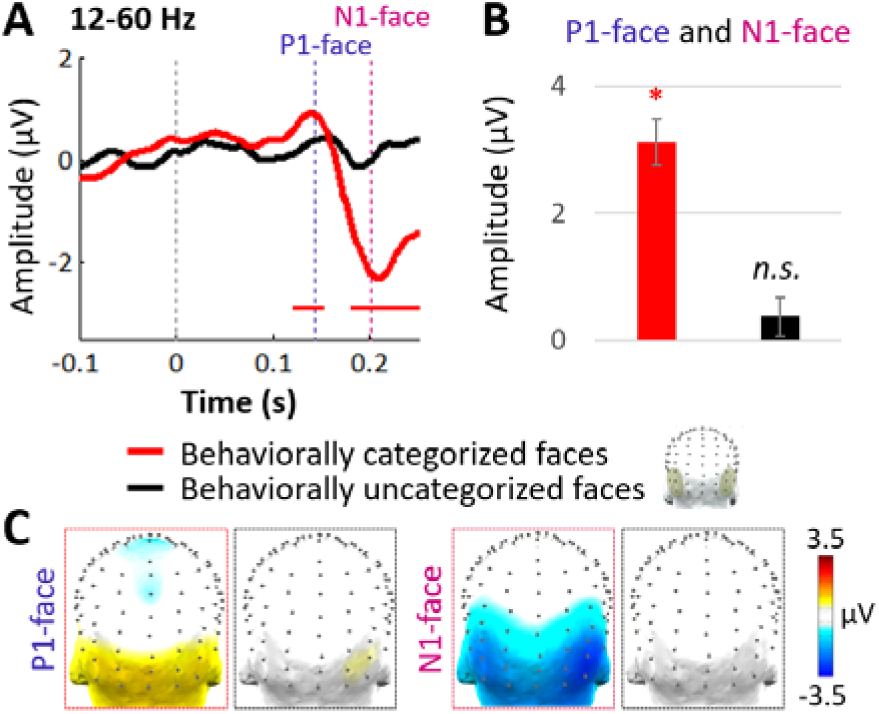
Time-domain responses (N = 16) across 12-60 Hz to all behaviorally categorized faces. **A**) Waveforms over the occipito-temporal ROI, to behaviorally categorized (in red) and uncategorized (in black) faces. Significant time points (p<.001) are indicated below for categorized faces (in red) and uncategorized faces (none). **B**) The amplitude across the P1- face and N1-face components, as defined in Experiment 1 (see Figure 6A), at the occipito-temporal ROI. Responses to behaviorally categorized faces (red) are significantly above zero, while those to behaviorally uncategorized faces (black) are not significant. **C**) Posterior topographies at the peak times of the P1-face and N1-face, for behaviorally categorized (red outline) and uncategorized (black outline) faces. Component times, taken from Experiment 1, are indicated in Panel A.

For statistical analyses, the amplitude of face-categorization responses was compared across the OT and MO ROIs (Table 2A). There was a significant main effect of *Region*, F_1,15_ = 19.5, p < .001, *η_p_*^2^ = 0.57, reflecting the larger amplitude for the OT (M = 0.94, SE = 0.12) than MO (M = 0.56, SE = 0.06) ROI in response to faces. There was also a main effect of *Frequency,* F_3.3,49_ = 5.94, p = .010, *η_p_*^2^= 0.86, with the low frequencies (12-3 Hz) being the highest and peaking at 12 Hz (M = 1.50, SE = 0.20), and then declining sharply as the frequencies continued to increase, with almost no amplitude at 120 and 60 Hz (M ≤ .02 µV). The interaction between these two factors, F_3.6,54_ = 10.2, p = .002, *η_p_*^2^= 0.91, was significant: most remarkably, there was no difference in amplitude across regions from 120-40 Hz (all differences < 0.02 µV), but a separation at 30 Hz and all lower frequencies (all differences > 0.32 µV).

For comparison, the stimulus-presentation responses were also evaluated over the OT and MO ROIs for statistical analyses (Table 2B). Here, the response was larger over the MO (M = 0.91, SE = 0.07) rather than OT (M = 0.63, SE = 0.04) ROI, leading to a significant main effect of *Region*, F_1,15_ = 43.8, p < .001, *η_p_*^2^= 0.75. The response also decreased from low to high frequencies, but with a different trend than for the face-selective response (compare Supplemental Figure 1 to Figure 2A). This produced a main effect of *Frequency,* F_1.5,22_ = 21.8, p < .001, *η_p_*^2^= 0.96. The interaction between these two factors, F = 21.8, p = .001, *η_p_*^2^= 0.92, was significant, as there was no amplitude difference across regions from 120-60 Hz (differences < 0.02 µV), but an increasing difference as the frequency decreased: at 40 Hz up slightly to 0.035 µV, at 30 Hz, 0.11 µV, and at all lower frequencies > .20 µV.

#### 2.2 Behavioral Responses (Experiment 2)

Behavioral face-categorization responses first emerged at 60 Hz, and accuracy increased steadily up to 68% at 20 Hz (50 ms; Table 3A). Beyond 20 Hz, accuracy neared ceiling between 12 to 3 Hz (83 to 333 ms; all accuracies above 95.6%). Overall, the differences in accuracy across frequency rates were highly significant, F_2.2,33_ = 217, p<.001, *η_p_*^2^= 0.94.

Response time (RT) showed less variation across presentation rates (Table 3B): from 3 to 24 Hz, there were no significant RT differences across conditions, F_4,60_ = 0.37, p = .83, *η_p_*^2^= 0.02, reflecting a stable face detection response at these rates. However, the response was slower at shorter presentation durations, and greatly delayed at 120 Hz (although note that only two participants had any (one or two) responses at this frequency). At the next shortest presentation rate, 60 Hz, 10 participants had some correct detection responses, contributing towards significant RT differences across frequency conditions from 3 to 60 Hz, F_1.5,12_ = 4.70, p = .040, *η_p_*^2^= 0.37. These differences persisted when removing 60 Hz, F_6,78_ = 6.84, p <.001, *η_p_*^2^= 0.35, in this case driven by slower responses at 40 Hz.

Inverse efficiency (IE) was also calculated, as a way to capture an overall measure of participants’ performance (Table 3C). This measure, as well as response accuracy, will be used in the following section to relate behavioral and EEG face-categorization responses.

### 3. The Relationship Between EEG and Behavioral Face-Categorization Responses

#### 3.1 The Correlation of Accuracy (Experiment 2) and Amplitude (Experiment 1)

The face categorization accuracy (Experiment 2) and EEG amplitudes (Experiment 1) seemed to show similar patterns across presentation rates (compare Panels B and C of Figure 1). Indeed, plotting the linear relationship between these two factors revealed a very strong correlation across all nine presentation rates, R^2^ = 0.96, p < .0001, as well as in the range where amplitude consistently decreased, 60-12 Hz, R^2^ = 0.97, p = .0002 (Figure 2C).

There are two possible interpretations of this observation: first, that the neural amplitude at higher presentation rates is decreased in response to each face to the same extent that the behavioral accuracy drops; or second, that the neural amplitude is decreased because some faces were not categorized at all, in the same way that the behavioral accuracy reflects the proportion of “hits” and “misses”. That is, either graded or all-or-none neural face categorization responses could explain this data. To fulfill our major goal of differentiating these alternative accounts, in the following sections we compare neural responses to behaviorally detected and undetected faces from Experiment 2.

#### 3.2 EEG Responses to Behaviorally Categorized or Uncategorized Faces (Experiment 2)

We examined non-periodic EEG responses in the time domain from Experiment 2 separately for 1) behaviorally categorized faces and 2) behaviorally uncategorized faces. Given that this experiment had relatively few trials per condition (about 30 on average, containing a mix of categorized and uncategorized faces), we combined responses across a range of presentation rates. Specifically, we used 12 to 60 Hz, in order to include a large number of trials, balanced in terms of the number of categorized (M = 62, SE = 7.1) and uncategorized (M = 60, SE = 5.7) faces across participants (after artifact rejection to remove blink-contaminated trials). Note that this approach is supported by neural face-categorization responses appearing qualitatively similar across presentation rates in Experiment 1 (see Section 4). Note also that these face-categorization responses were confounded with behavioral detection (pre-)motor activity, with response times occurring on average at 535 ms over these presentation rates (SE = 15.1 ms). We thus focused the analyses on the first two face-selective components, the P1-face (onset at about 100 ms, 144-ms peak latency) and N1- face (202-ms peak latency), as defined in Experiment 1 (Section 4.3; and reported in previous studies with this paradigm, see e.g., Rossion et al., 2015; Retter & Rossion, 2016a; Quek & Rossion, 2017). Finally, note that although faces appeared periodically in Experiment 1 and non-periodically in Experiment 2, comparing neural face categorization responses across experiments is valid because such responses are immune to the periodicity of the stimulus (Quek & Rossion, 2017).

For behaviorally categorized faces, significant neural responses (event related potentials) were present during the windows of the P1-face and N1-face; for uncategorized faces, little to no responses were apparent, exhibiting no significance (Figure 3A). Examination of the P1-face and N1-face peak-time deflections together over the bilateral OT ROI also showed a significant response to behaviorally categorized faces (M = 3.12 µV, SE = 0.37 µV), t_15_ = 8.47, p <.001, d = 3.0, but an insignificant response to behaviorally uncategorized faces (M = 0.38 µV, SE = 0.30 µV), t_15_ = 1.24, p = .12, d = 0.44 (Figure 3B). The topographies of these face-selective components for behaviorally categorized faces were maximal over the OT region; little activation was observed across the scalp to behaviorally uncategorized faces (Figure 3C).

#### 3.3 Behavioral Face Categorization Predicts EEG Face-Categorization Amplitude (Experiment 2)

All-or-none neural responses, as reported in the previous section, could thus explain the gradient of response amplitudes reported in Experiment 1: this gradient reflects the proportion of absent neural responses to uncategorized faces and full neural responses to categorized faces. To further test this hypothesis, we examined whether the neural face-categorization amplitude would be reduced at higher relative to lower rates in the *same proportion* that the behavioral face-categorization accuracy was reduced.

To this end, we examined the neural responses to non-periodically presented faces in Experiment 2 in the time domain, for behaviorally categorized and uncategorized faces together. Specifically, we examined neural responses across two relatively high presentation rates (24 and 30 Hz) with fewer correct detections, and compared them to two relatively low presentation rates (12 and 20 Hz) with more correct detections, for nine participants meeting a minimum behavioral accuracy threshold (see Methods). Again, we combined responses across multiple rates given the limits from the number of face presentations in this experiment. As before, we constrained the analysis to the first two face-selective components.

The amplitude of the neural responses to behaviorally categorized and uncategorized faces together appeared reduced at 24-30 Hz, relative to 12-20 Hz, at the times of the P1-face and N1-face; in contrast, the response to behaviorally categorized faces only did not appear to differ across presentation rates (Figure 4A). This was evident over the bilateral occipito-temporal ROI, which appropriately captured face-categorization responses for both conditions (Figure 4B).

**Figure 4.**
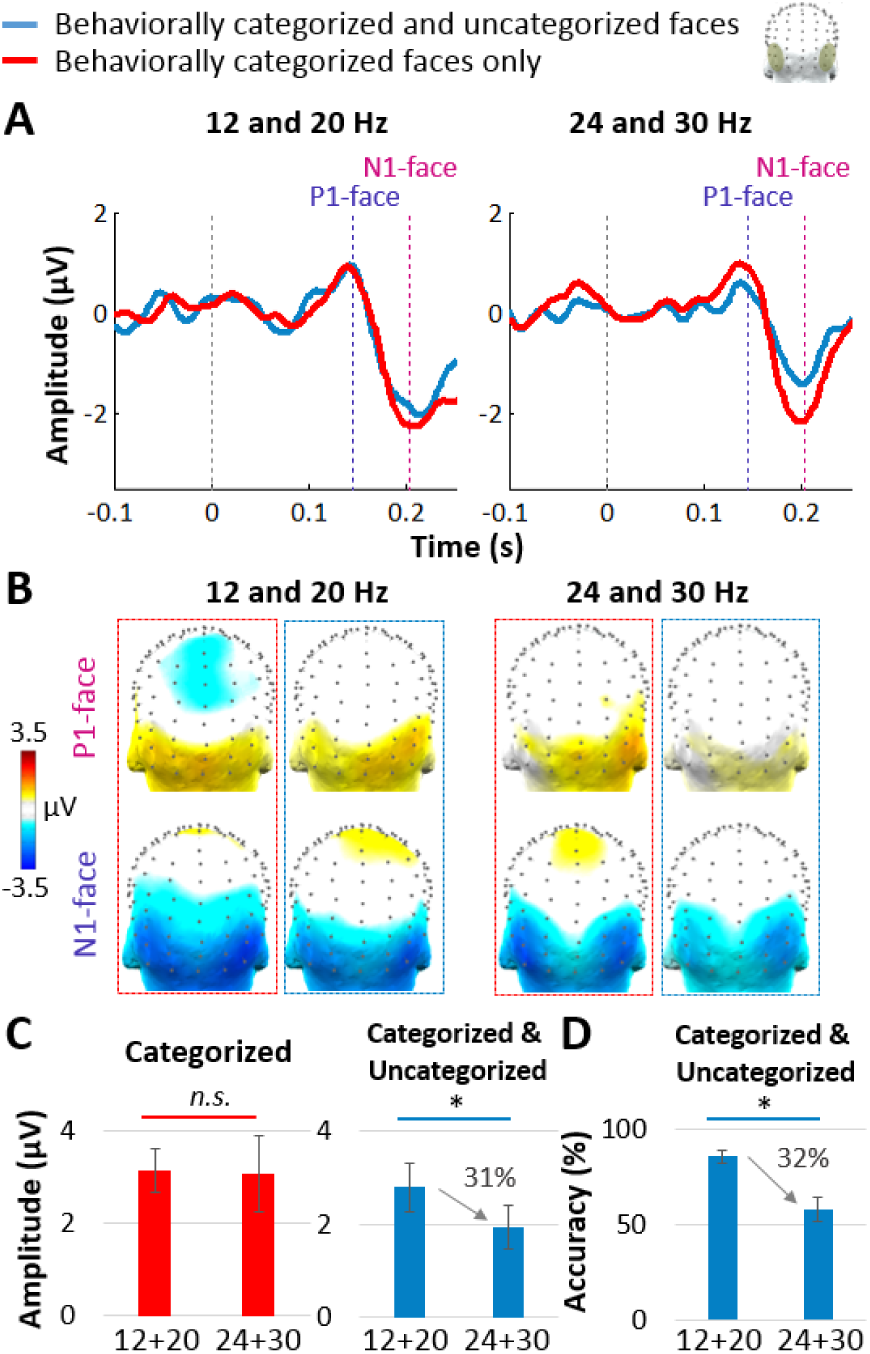
Time-domain responses from Experiment 2 to behaviorally categorized and uncategorized faces together (blue), compared to categorized faces alone (red). There were fewer categorized faces at the higher (24 and 30 Hz), relative to lower (12 and 20 Hz) frequencies; results are shown for participants who had a minimum of 20 artifact-free face-categorizations at both frequency pairs (N = 9). **A**) Waveforms of the bilateral OT ROI for 12 and 20 Hz (left), and 24 and 30 Hz (right), for both behaviorally categorized faces only (red) and behaviorally categorized and uncategorized faces combined (blue). The indicated times of the P1-face (144 ms) and N1-face (202 ms) components are taken from Experiment 1 (see Figure 6A). **B**) Scalp topographies of these two components; outlines indicating condition are colored as in the previous panels. **C**) The combined amplitude of the P1-face and N1-face components, at the OT ROI. Responses to behaviorally categorized faces alone did not differ across these conditions (left), but responses to behaviorally categorized and uncategorized faces together were significantly larger at 12 and 20 Hz than at 24 and 30 Hz (right). **D**) The percent accuracy dropped from 12 and 20 Hz to 24 and 30 Hz by about 32%; note the correspondence with the 31% decreased EEG amplitude across these conditions in Panel C for to behaviorally categorized and uncategorized faces.

When behaviorally categorized faces were considered alone, a significant difference in the combined amplitude of the P1-face and N1-face components was not found across 12 and 20 Hz, (M = 3.12 µV; SE = 0.47 µV) vs. 24 and 30 Hz (M = 3.38 µV; SE = 0.72 µV), t_8_ = −0.06, p = .48, d = 0.02, replicating the finding in the previous section (Figure 4C, left panel). However, the combined amplitude was significantly larger at 12 and 20 Hz (M = 2.80 µV; SE = 0.53 µV) than at 24 and 30 Hz (M = 1.95 µV; SE = 0.48 µV) when behaviorally categorized and uncategorized faces were considered together, t_8_ = 2.29, p = .026, d = 0.56 (Figure 4C, middle panel).

Behaviorally, the accuracy was reduced by 32.2% from 12 and 20 Hz to 24 and 30 Hz (from 85.8%, SE = 3.27%, to 58.1%, SE = 6.44%) for these participants, t_8_ = 6.35, p = .0001, d = 1.80 (Figure 4D). The EEG amplitude across the P1-face and N1-face components was decreased similarly, by 30.6%, from 12 and 20 Hz to 24 and 30 Hz (from 2.80 µV to 1.95 µV), when responses to behaviorally categorized and uncategorized faces were combined. This corresponding decrease of behavioral accuracy and amplitude supports the hypothesis that missed face categorizations contribute proportionally to decreased neural response amplitudes at shorter presentation durations.

### 4. Qualitative Investigation of Face-Categorization Responses

#### 4.1 Multi-Harmonic Response Distributions (Experiment 1)

In the temporal frequency-domain amplitude spectrum, face-categorization responses were distributed across 1 Hz and its harmonics below 20 Hz (Figure 5A). Harmonic responses were generally higher at lower frequencies, and strongest between 1 and 10 Hz, with a local peak typically around 6 Hz. Stimulus-presentation responses, which occurred at higher frequencies, were spread across fewer harmonics (Figure 5C). In contrast to face-categorization harmonic responses, they were typically dominated by the amplitude of the first harmonic frequency and decayed exponentially as harmonic frequency increased.

**Figure 5.**
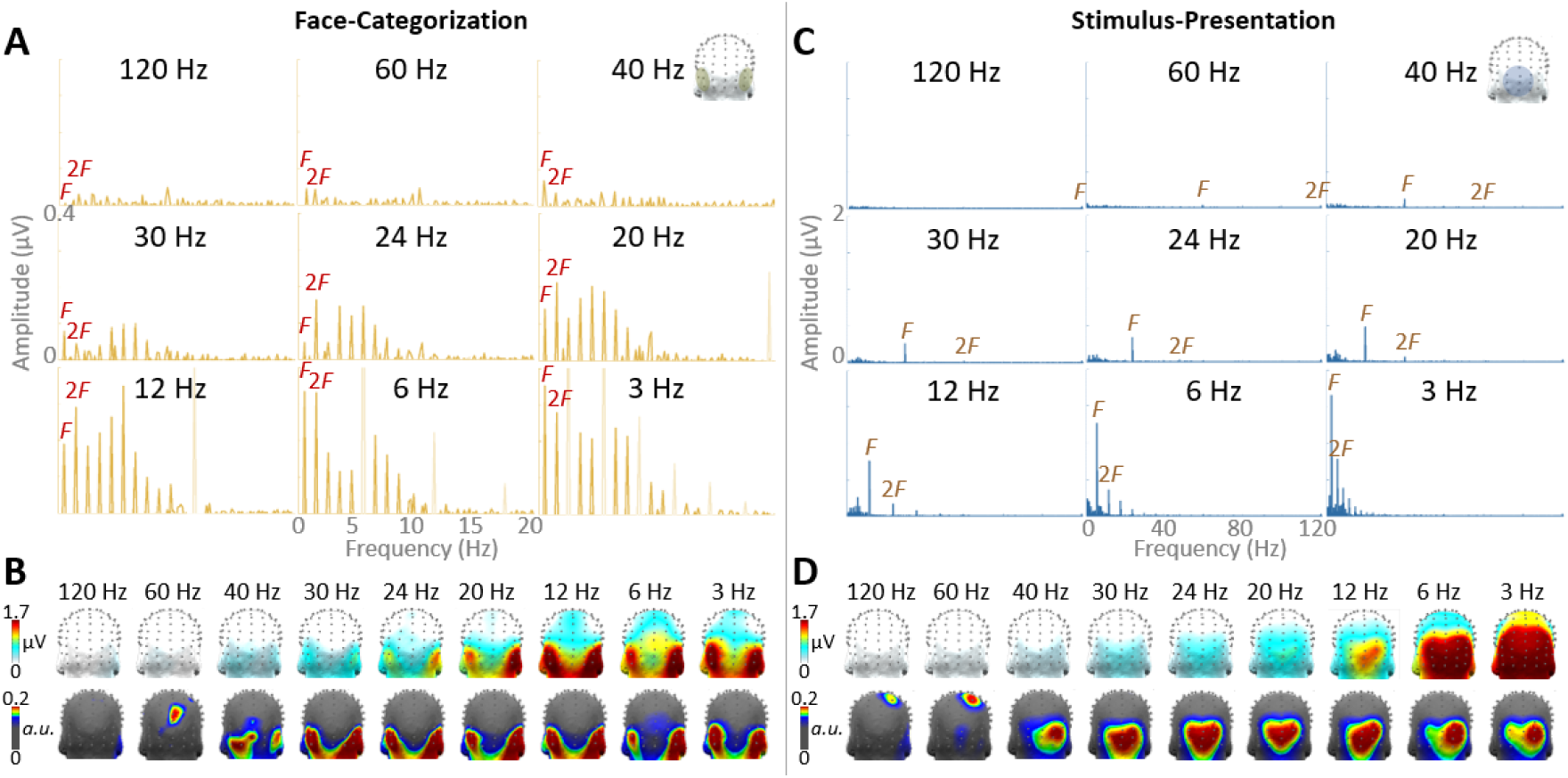
Face-categorization responses characterized in the frequency domain, and compared to stimulus-presentation responses. **A**) Face-categorization baseline-subtracted amplitude spectra. Face-selective responses were tagged at F (= 1 Hz) in every condition, with higher harmonics occurring at 2F (2 Hz), 3F (3 Hz), etc. (the first two harmonics are labeled above the spectra). The data are shown over the region of the scalp (bilateral occipito-temporal ROI) and frequency range (up to 20 Hz) that were selected to capture face-selective responses. Higher harmonic frequencies coinciding with stimulus-presentation are drawn in very light yellow, and may surpass the plotted amplitude range. For ease of comparison across conditions, all graphs are plotted with common axes. **B**) Face-categorization response topographies. Top row: grand-averaged baseline-subtracted summed-harmonic topographies. Bottom row: the same responses normalized. **C**) Stimulus-presentation responses are tagged at the frequency of presentation (F) corresponding to the condition name. As in Panel A, the first two harmonic frequencies are labeled above the spectra for each condition. The data is shown over the region of the scalp (medial-occipital ROI) and frequency range (up to 120 Hz) that were used to capture stimulus-presentation responses. All graphs are plotted with common axes, although different from those in Panel A. **D**) Stimulus-presentation topographies. Top row: grand-averaged baseline-subtracted summed-harmonic topographies. Bottom row: the same responses normalized.

Note that in the previous sections, given the ranges of the distributed harmonic responses, as well as expected ranges from previous studies (e.g., Jacques, Retter & Rossion, 2016; Retter & Rossion, 2016a; Retter et al., 2018), baseline-subtracted harmonics were summed from 1 to 20 Hz for face categorization responses (Retter & Rossion, 2016a). For stimulus presentation responses, in order to encapsulate the full range of presentation frequencies used across conditions, baseline-subtracted harmonic responses were summed up to 120 Hz (see Methods). The face-categorization amplitudes across presentation rates were shown in Figure 2A, and can be compared to the corresponding stimulus-presentation amplitudes in Supplemental Figure 1.

#### 4.2 Response Topographies (Experiment 1)

The a-priori bilateral occipito-temporal (OT) region of interest was verified for its ability represent face-categorization responses. Indeed, the OT ROI represented 7 to 9 out of the 10 channels displaying the maximal face-categorization response amplitudes across presentation conditions from 3 to 40 Hz, although only 2 at 60 Hz and 0 at 120 Hz (Figure 5B: top row). For further verification, we controlled for differences in response amplitude across presentation duration conditions by normalizing the response topographies: this shows occipito-temporally dominated responses again from about 40 to 3 Hz (Figure 5B: bottom row). Note that for response quantification, this region was also broken into the right and left hemispheres to assess potential right lateralization effects generally predicted for face perception.

In terms of lateralization, the ROT appeared to consistently have a higher amplitude than the LOT, except at 3 Hz. Indeed, the lateralization index (see Methods) revealed a right lateralization (values > 0) across all significant conditions (30–3 Hz), except 3 Hz (M = −3.13, SE = 4.33; see Figure 3A for data at each the right and left OT ROI). The right lateralization was typically low, ranging from 2.18 to 16.1 across 30 to 6 Hz; it was maximal at 20 Hz (SE = 11.6). Yet, at the group level, there were no significant differences in lateralization across conditions, F_1.4,11_ = 1.35, p = .31, *η_p_*^2^= 0.38.

The responses to stimulus presentation were shown to contrast with the face-categorization responses. These stimulus-presentation responses typically occurred maximally over the medial occipital (MO) scalp region (Figure 5D: top row). Their response topographies varied somewhat as the presentation frequency varied, being most different at 60 and 120 Hz, where a centro-parietal region dominated the responses (Figure 5D: top row).

#### 4.3 Spatio-Temporal Dynamics (Experiment 1)

The face-categorization responses from the previous section were re-examined in the time domain to explore potential qualitative differences across presentation rates. The following analyses use data selectively filtering out the stimulus-presentation frequency (see Methods; for unfiltered data, see Supplemental Figure 2). Four prominent time-windows of interest appeared across conditions, corresponding to those reported previously as P1-face, N1-face, P2-face, and P3-face (at a 12.5-Hz stimulus-presentation rate; Retter & Rossion, 2016a). These time-windows encompassed nearly all significant deviations over the bilateral OT ROI (Figure 6A). Each of these deflections, or “components”, was significant across three to six frequency conditions, without any clear trend of component differences as stimulus presentation duration varied. Across significant conditions, these four components peaked over the OT ROI at: 144 ms (SE = 1.95 ms); 202 ms (SE = 5.09 ms); 281 ms (SE = 5.34 ms); and 463 ms (SE = 12.8 ms) post face-stimulus onset.

**Figure 6.**
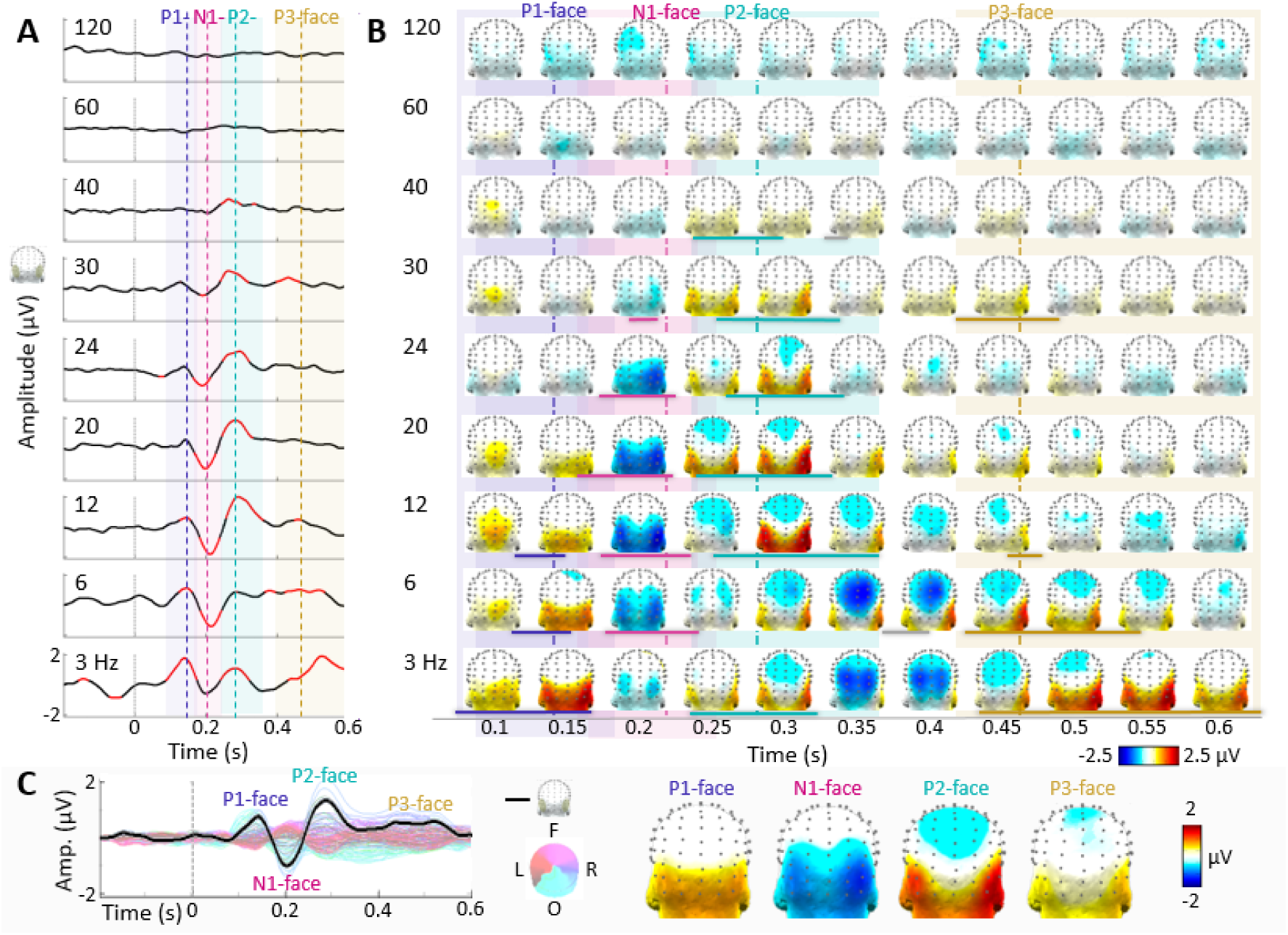
Time-domain responses to face presentation, with each frequency condition filtered to remove the response at its respective stimulus-presentation frequency. Four components were present repeatedly across conditions: P1-face, N1-face, P2-face, and P3-face, and are indicated by a vertical line showing the average peak time, and shading depicting the range, across significant conditions. **A**) Waveforms of the bilateral OT ROI in response to face stimulus onset (0 s). For each condition, significant time periods (p<.001) are plotted in red, while those insignificant are plotted in black. **B**) Scalp topographies, plotted from 0.1-0.6 s post-stimulus onset, every 0.05 s. Significant response periods are underlined in the color of their respective component for each condition; significant periods not corresponding to one of these four components are indicated in gray. **C**) An average of all conditions with significant face-categorization responses in the frequency domain. The bilateral occipito-temporal ROI is plotted in thick black, superimposed above the data from all 128 EEG channels, colored accordingly to the adjacent 2D scalp topography (F: frontal; R: right; O: occipital; L: left). The topographies are plotted to the right at the peaks of the P1-face (142 ms), N1-face (202 ms), P2-face (286 ms), and P3-face (519 ms) for this waveform.

These deflections, and responses more generally across time, occurred mainly over the bilateral OT ROI, with some coverage of the occipito-inferior, occipito-parietal, and medial-occipital regions (Figure 6B). In concurrence with the frequency-domain analysis, no significant deflections were found for the conditions at 120 or 60 Hz; however, one significant time-window was found for 40 Hz at the P2-face time, perhaps contributing to the non-significant amplitude in the frequency domain over the OT region for this condition (see again Figure 4B). To characterize a frequency-independent face-categorization response, an average of all conditions with significant face-categorization responses as determined from the frequency domain (3–30 Hz) was created (Figure 6C). Over the OT ROI, this response first reached significance at 103 ms post-stimulus onset, and retained significance for 460 ms: it was consistently significant at the times of the P1-face (103 to 159 ms), N1-face (175 to 234 ms), P2- and P3-face (247 to 560 ms).

In addition to the four main time-windows, a significant time period emerged in the 6 Hz condition between 366–398 ms, peaking at 380 ms. This 6-Hz OT positive deflection was accompanied by a large medial parieto-occipital negativity, peaking about 15 ms earlier, which was previously described as the “N2-face” (with a 5.88-Hz sinewave presentation; Jacques, Retter & Rossion, 2016). This response signature is also apparent when stimuli are presented at 3 Hz, and perhaps at 12 Hz and 24 Hz, although not reaching significance over the OT scalp ROI in these conditions (see again Figure 6B).

#### 4.4 Summary

In sum, face-categorization responses were qualitatively similar in terms of their harmonic response distributions in the frequency domain, their summed-harmonic topographies, and their spatio-temporal dynamics.

### 5. Individual Differences in Generic Face Categorization

#### 5.1 The Diagnosticity of Presentation Rate (Experiments 1 and 2)

Here, we aimed to investigate whether individuals’ EEG amplitudes (Experiment 1) were related to their behavioral face-categorization performance (Experiment 2) overall. As a first step, we checked what effect presentation rate had on the range of individual differences, both behaviorally and neurally. Individual participants were ranked for their behavioral face-categorization performance based on their inverse efficiency (IE) score across all presentation rates (range = 0.65 to 1.36; M = 0.97; SE = 0.054; see Table 3C).

The best participant at behavioral face detection (S07) was contrasted to the worst (S01), in terms of percent accuracy and EEG amplitude. (Note that the same participants would have been ranked as the best and worst if accuracy alone was used as a metric.) Large differences between these participants in both behavioral and EEG amplitude responses were apparent in the mid-frequency range, from 20–30 Hz (50–33.3 ms; Figure 7A). No differences were apparent across these participants at the lowest three presentation rates (3–12 Hz, i.e., 333–83.3 ms) in either measure, and differences were greatly reduced by 40 and 60 Hz (25 and 16.7 ms, respectively) until they were abolished at 120 Hz (8.33 ms). The data of all participants, grouped into quartiles, followed these trends, at least between the best 25% and worst 25% of participants (Figure 7B).

**Figure 7.**
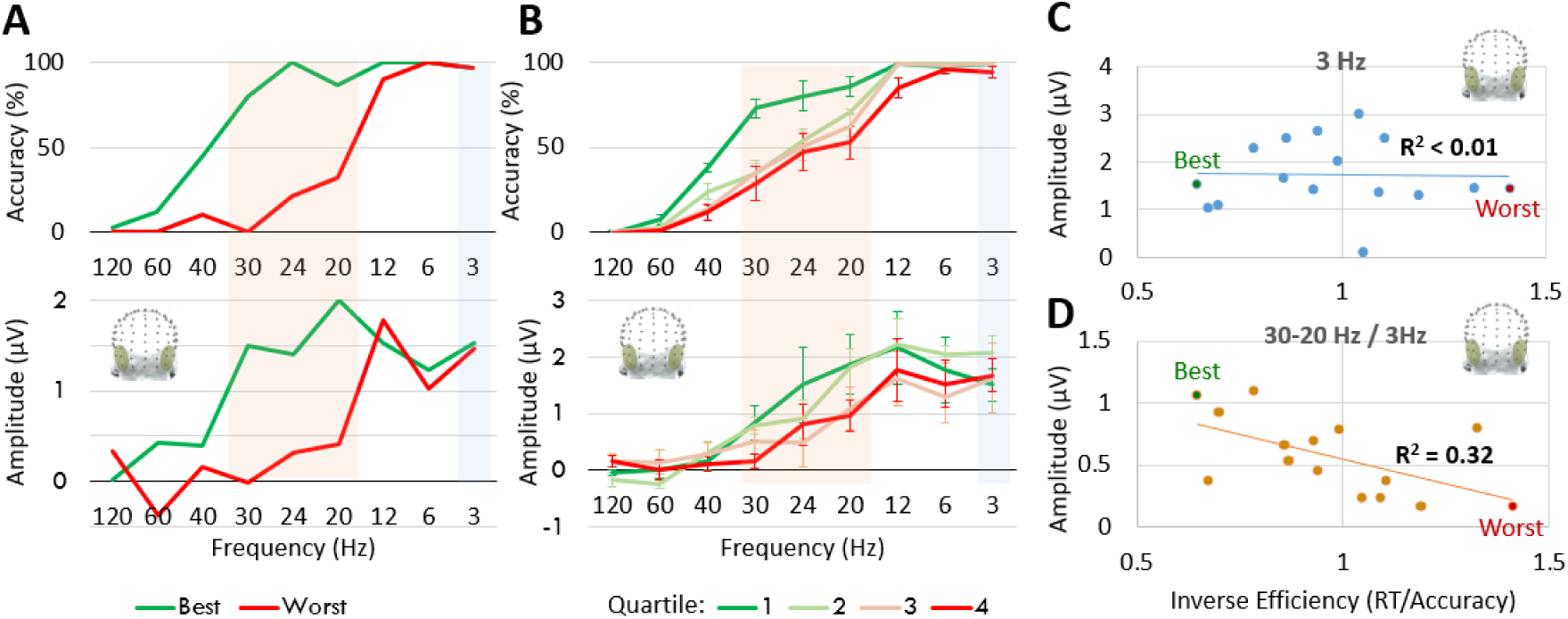
Individual differences in behavioral performance related to EEG amplitude for face categorization. **A**) Presentation rates diagnostic of individual differences. A comparison of the participant with the best vs. worst behavioral (inverse efficiency; IE) responses across presentation rates. The highlighted sections indicate data ranges used in panels C and D. Above: Accuracy; Below: Summed baseline-subtracted harmonic amplitude over the bilateral OT ROI. **B**) As in Part A, except the 16 participants were split into quartiles of four participants each, ranked 1^st^-4^th^ behaviorally (Group 1), 5^th^-8^th^ (Group 2), 9^th^-12^th^ (Group 3), and 13^th^-16^th^ (Group 4). **C)** The non-significant correlation of individuals’ IE (across all presentation rates) with their OT ROI EEG amplitudes at 3 Hz (highlighted in blue in the previous two panels). **D)** The significant correlation between individuals’ mid-frequency OT EEG amplitudes (averaged over 20, 24, and 30 Hz, highlighted in orange in the previous two panels), weighted by the amplitude at 3 Hz. In both panels C and D, the data from the participants yielding the best and worst behavioral performance are labeled, respectively.

The lack of significant EEG responses in the high presentation rates (40–120 Hz) would prevent a relationship between individuals’ EEG and behavior at this range. At low frequencies (3–12 Hz), behavior near ceiling would prevent a relationship between individuals’ EEG and behavior. The best separation in EEG response amplitudes was produced at 20 Hz, and behavioral responses seemed well-separated across the mid-frequency range (from 20–30 Hz; 50–33.3 ms). In the following section, therefore, we will relate individuals’ EEG response amplitudes to their behavioral performance using this mid-frequency range.

#### 5.2 The Correlation of Individual Participant’s EEG Amplitudes (Experiment 1) and Behavioral Accuracy (Experiment 2)

To examine whether there was an inherent relationship between individuals’ behavioral face-categorization performance (IE scores, computed across presentation rates) and EEG face-categorization amplitudes, we correlated these measures at the slowest presentation rate, 3 Hz (333 ms). This correlation was not significant, without any apparent trend, R^2^ = .0006; p = 0.93 (Figure 7C), which is likely explained by variable physiological factors influencing the EEG response across participants (see Discussion).

When correlating individuals’ behavior and amplitudes in the mid-frequency range, from 20 to 30 Hz, as defined in the previous section, the correlation was improved but still not significant, R^2^ = 0.22; p = .067. However, when weighting the individuals’ mid-frequency range EEG amplitudes by their baseline amplitudes at 3 Hz, a significant correlation was found, R^2^ = 0.32, p = .043 (Figure 9D). This measurement is reflective of the change in amplitude of the EEG response from its amplitude at a presentation rate with ceiling-level behavioral face-categorization performance, and thus likely reflects the percent change in behavioral performance as presentation rate decreases. Yet, note that the weighted correlation was not significantly stronger than the unweighted correlation, Z = 0.99, p = .16 (one-tailed).

## Discussion

With a novel sweep frequency-tagging visual categorization paradigm, we (1) determined the minimal and optimal presentation duration necessary to elicit both behavioral and neural face-categorization responses in human observers, (2) related these responses at the group and individual levels, and (3) determined whether visual categorization is graded or all-or-none. In a first experiment, we presented faces periodically within non-face object RSVP sequences, and investigated implicit response in the EEG frequency domain. In a second experiment, we presented faces non-periodically within non-face object RSVP sequences, and investigated explicit behavioral categorization responses, as well as their associated electrophysiological responses.

### All-or-None Categorization Responses

Our results suggest that stimuli are either categorized as faces and elicit full responses, even at very brief presentation durations (e.g., 17 ms), or they are not categorized, even sometimes at relatively low presentation durations (e.g., 50 ms), and no face-selective responses are elicited. Indeed, no significant neural responses to behaviorally uncategorized faces were present in the categorical, or differential, neural responses of Experiment 2. However, across presentation rates from 12 to 60 Hz (83.3 to 17.6 ms), large deflections were present at the times of the P1-face and N1-face in response only to behaviorally categorized faces (Figure 3). Additionally, these face-categorization responses obtained at 12 and 20 Hz were no larger than at 24 and 30 Hz when only behaviorally categorized faces were taken into account (Figure 4).

All-or-none categorization responses are not inconsistent with the apparent gradient of response amplitude observed as presentation rate changed in Experiment 1 (Figure 2A). Instead, we propose that the proportion of categorized to uncategorized faces drives graded amplitude differences, similar to how behavioral categorization of “hits” and “misses” drives graded behavioral response accuracies. Suggesting such an explanation, the behavioral and frequency-tagged neural face-categorization responses were strongly correlated across Experiments 2 and 1, respectively (Figure 2C). Moreover, the decrease in the percent of faces that were categorized behaviorally (a drop of 31% when comparing 24 and 30 Hz to 12 and 20 Hz) matched the decrease in the amplitude (32%) of the neural face-categorization response as quantified in the same experiment (Figure 4C and D).

It should be noted that the correspondence of neural and behavioral measures as image processing time varies (as a function of presentation duration and/or backward-mask latency) is a replication of a number of previous studies (e.g., Rolls & Tovee, 1994; Kovács, Vogels & Orban, 1995; Vanni et al., 1996; Grill-Spector et al., 2000; Keysers et al., 2001; Bacon-Macé et al., 2005). Note that such studies, if interpreted in this light at all, have been taken as evidence for graded perceptual effects, attributing the decrease in response as a decrease in the response to each stimulus presented (e.g., see Keysers et al., 2001; Bacon-Mace et al., 2005). Our account and evidence of this relationship as variations in the *proportion* of all-or-none neural responses, is to our knowledge novel and suggests a different underlying process.

Further in line with all-or-none categorization, even as neural amplitude varied with presentation rate in Experiment 1, face-categorization responses remained qualitatively similar in terms of scalp topography (consistent maximal responses over the right occipto-temporal region: Figure 5B), time course (no trends by presentation speed affecting the P1- face, N1-face, P2-face, or P3-face deflections: Figure 6), and the distribution of amplitude across face-categorization harmonic frequencies (Figure 5A). Qualitatively similar face-categorization responses in both the frequency and time domain have also been reported with this frequency-tagging categorization paradigm across studies using different stimulus-presentation rates from about 6 to 20 Hz (e.g., Rossion et al., 2015; Retter & Rossion, 2016a; Quek & Rossion, 2017; Retter et al., 2018) and across variations of stimulus visibility (i.e., low-pass filtering, Quek et al., 2018). Moreover, raw responses to faces or other visual objects appear similar both quantitatively and qualitatively when using variable image presentation durations, for example from about 40 to 500 ms (Cichy, Pantazis & Oliva, 2014).

While many arguments have been made for a progressively increasing response amplitude enabling stimulus recognition in the visual cortex (e.g., Kleinschmidt et al., 1998; Bar et al., 2001; Moutoussis & Zeki, 2002; Jemel et al., 2003), all-or-none neural responses have been proposed previously for stimulus detection or identification (e.g., Del Cul, Baillet & Dehaene, 2007; Quiroga et al., 2008; see also Navajas et al., 2013; Sekar, Findley & Llinas, 2012; Sekar et al., 2013 for unmasked stimuli). However, the stimuli used in these previous studies are limited in range and well-segmented (e.g., a few letters, numbers or words in Del Cul et al., 2007; Sekar, Findley & Llinas, 2012; Sekar et al., 2013; a few repeated exemplar images in Quiroga et al., 2008; full front segmented faces in Navajas et al., 2013). Moreover, the timing of these all-or-none neural responses is unclear, with relatively late all-or-none responses often reported (e.g., more than 270 ms in widespread regions in Del Cul et al., 2007; more than 300 ms in Quiroga et al., 2008 for neurons in the human medial temporal lobe). Compared to these studies, our study provides original evidence for all-or-none neural categorization responses, given that: 1) it directly measures categorization (i.e., a differential and generalizable response across widely variable exemplars); 2) through frequency tagging, the neural response is objectively identified and quantified relative to surrounding noise level in the frequency domain; 3) the neural markers of face categorization in this paradigm are known to originate from face-selective neuronal populations in the ventral occipito-temporal cortex (see Jonas et al., 2016; Gao et al., 2018); and 4) it shows unambiguous face categorization responses emerging at a 17 ms duration/SOA, and with a latency of 100 ms following stimulus onset.

Finally, the interpretation of all-or-none categorization responses does not imply that at very short presentation durations the perception of a face stimulus is full, i.e., that all the potential visual information, such as age, gender, and identity, has been extracted from the image. Thus, perception may be graded in the sense that a face may be recognized as a face before its identity is recognized. However, we suggest that the categorization process itself is all-or-none: if neural responses differentiating faces from objects and generalizing across face exemplars are elicited at all, they are elicited fully.

### Why Are Some Stimuli Not Categorized?

While high-level (i.e., category-selective) visual processes are all-or-none, they may still be reliant on the gradual accumulation of low-level visual information in early visual areas of the brain (e.g., Windey et al., 2014). At an early stage, we theorize that if enough visual information is assembled from the stimulus within about 20 ms, it goes on to develop and trigger a full category-selective response after approximately 100-ms post-stimulus onset (e.g., Crouzet et al., 2010; Crouzet & Thorpe, 2011; Liu et al., 2009; Jacques et al., 2016; Retter & Rossion, 2016a; see Carandini et al., 2005 for a review of this process up to V1). Indeed, in previous studies, the minimum duration to evoke a neural response was between about 12-20 ms when images were backward-masked (e.g., with scrambled images; see Breitmeyer, 1984); this measure was replicated with a wide variety of recording techniques, including single unit in monkeys (Rolls & Tovee, 1994; Kovács, Vogels & Orban, 1995; Keysers et al., 2001), MEG (Vanni et al., 1996), fMRI (Grill-Spector et al., 2000), and EEG (Bacon-Macé et al., 2005) in humans. Here, a highly sensitive 20 ms window is consistent with responses occurring rarely when interrupted with a backward-mask after 17 ms, and large increases in the amount of categorization with the next durations (SOAs) of 25 and 30 ms.

However, if early visual processing is interrupted after about 20 ms but before the category-selective response onset (at approximately 100 ms here, and also in Retter & Rossion, 2016a), perceptual categorization may still not occur. Beyond its initiation, visual information persists in the visual system, e.g., for about 60-100 ms, in early (e.g., V1) brain areas (e.g., see the Discussions of Rolls & Tovee, 1994; Keysers et al., 2001). At these stages, the accumulation of visual information may still be disrupted from masking producing competition in early visual areas (see the “neural competition” theory of Keysers and Perrett, 2002; see also the discussion of Potter, 2012). In these experiments, the earlier a mask is presented following a face, the shorter the sensory processing time, and the more likely that the face will not be categorized. In our experiment, missed face categorizations occurred more often at shorter presentation durations, and even up to SOAs of 50 ms (20 Hz; Figure 2). At each face presentation, whether or not the image is categorized or not likely depends on many factors, including: properties of the face image (e.g., how central or high-contrast the face is, or the face’s head orientation), the overlap of low-level properties with its flanking images (Crouzet & Thorpe, 2011; see again Potter, 2012; but also Maguire & Howe, 2016; Broers et al., 2018), the local context (e.g., whether the preceding face has been categorized), and the neural representation of faces in the stimulated brain.

Note that in this case, a strong effect of masking is not predicted in high-level visual areas, because stimuli temporally flanking the faces consist of non-face objects for at least 1 s before and after (in contrast, see Retter & Rossion, 2016a for a case of high-level face competition when face stimuli are presented with less than about 400 ms SOAs). In previous forward- and backward-masking studies, early competition from masking stimuli was shown to contribute to reduce mean response amplitude when target stimuli were shown for as low as 28 ms; in contrast, with a “gap” between stimuli resulting in SOAs of at least 100 ms, a reduction in amplitude was not produced (Keysers et al., 2005; Retter et al., 2018). Nevertheless, we suggest that if a face is not categorized because of low-level interference before about 100 ms, it does not enter perceptual awareness (see also Edelman, 1978, pp. 81- 88).

### The Minimal Speed for Conscious Face Categorization

A minimum duration of 17 ms (60 Hz presentation rate) was required to elicit occasional face categorization responses in both neural and behavioral measures. This minimal duration is conservative in that it results from highly variable images: the faces varied in size, eccentricity, head orientation, gaze direction, expression, etc. Nevertheless, behavioral face categorization responses were present significantly, but minimally, with 17 ms of image presentation (60 Hz; M = 3.03%, SE = 1.26%, with more frequent responses than false alarms evident at this rate in half (8/16) of the individual participants). In contrast, face categorization was clearly not possible at the shortest presentation duration tested, 8 ms (120 Hz), at which only one participant had one categorization, albeit with no false alarms.

At a neural level, face-categorization responses occurred occasionally at 17 ms over the bilateral occipito-temporal ROI (evident in Experiment 2; data not shown), although these were significant for only one participant in Experiment 1, who had the highest accuracy at this presentation rate. Neural responses were instead not significant at 17 ms for most participants in Experiment 1, likely due to an insufficient number and proportion of detected trials to separate EEG signal from noise: e.g., a participant detecting the average 3% of faces in the second experiment would only have detected about 4 of 126 faces presented in the first experiment at 17 ms. In comparison, the best participant behaviorally detected 12.5% of faces in the second experiment, predicting about 16 faces detected in the first experiment at 17 ms. Note that these 16 responses to faces presumably compete for amplitude with the 110 faces that were not detected, such that recording any significant response is notable, and likely due to the high signal-to-noise ratio afforded from frequency-domain analyses of periodic responses (Norcia et al., 2015).

Perhaps due to such factors, group-level significance was not evident here until 33 ms, when group mean accuracy was 22% (predicting about 28/126 faces detected in the first experiment; Figure 1; Table 1). Across participants, the minimum presentation duration required for a significant EEG face-categorization response over the bilateral OT ROI ranged from 17 to 83 ms (Figure 1). Note that while a linear relation between behavior and neural responses was significant across all presentation rates here, a threshold for recording neural responses predicts a non-linear relationship between behavior and neural responses at the fastest presentation rates (i.e., below about 17 ms here; see Figure 2C).

Although very short, the minimum duration of 17 ms for face categorization is consistent with the results of several previous studies, both at behavioral and neural levels. Notably, in a study by Keysers and colleagues (2001; 2005; see also Perrett et al., 2009), the temporal tuning of the responses of single cells and populations in macaque monkeys was investigated in response to monkey heads presented at varying orientations in RSVP sequences. Neural firing rates were shown to decrease at faster presentation rates, but responses to non-periodic target stimuli could still be detected at up to 14 ms (with 300-400 trials), the fastest rate in that experiment. Behavioral accuracy was measured separately with human participants, which also decreased with presentation speed and remained significant up to 14 ms. A later behavioral study using RSVP closely replicated this presentation duration limit for high-level visual perception: Potter et al. (2014) showed recognition of various natural images (e.g., people or flowers) by human participants at 13 ms, the fastest rate in their experiment. Further, additional studies have confirmed the presence of face (and object) detection responses at 16-17 ms with behavior, ECoG, and MEG, in masked or short RSVP sequences (Grill-Spector & Kanwisher, 2005; Fisch et al., 2009; Mohsenzadeh et al., 2018). The common limit of approximately 15 ms across these studies perhaps reflects something general in visual perception: the minimum amount of time required for a visual stimulus to be coherently processed in early visual areas, before triggering a high-level classification response.

It should be noted that our study was limited in its ability to present stimuli at many durations near potential thresholds, due to the constraints of monitor refresh rates. For example, durations between 17 and 8 ms could not be tested, since these durations consist of two and one screen display frames, respectively, on a 120 Hz monitor. This limitation in available presentation durations has similarly affected many previous studies, and for example limits pinpointing whether a limit is at 13 (Potter et al., 2014), 14 (Keysers et al., 2001), 16 (Fisch et al., 2009), or 17 ms (Grill-Spector & Kanwisher, 2005; Mohsenzadeh et al., 2018). Accordingly, it would also be difficult to assess subtler questions, such as whether faces may be identified faster than other objects around such upper limits (after 17 ms, three display frames at 120 Hz determined the next available duration at 25 ms; but see Fisch et al., 2009; and for a different approach, see Rousselet et al., 2003). Such questions require faster framerates than most current displays afford.

### The Optimal Speed of Face-Categorization

The maximal behavioral accuracy and neural amplitude occurred at the same time, around 80 ms (12 Hz; Figure 2). This optimum time aligns well with ranges from previous studies: for example, maximum accuracy and EEG responses were reported at 81 ms for categorizing animal images followed by dynamic visual masks (adjacent durations of 44 and 106 ms; Bacon-Mace et al., 2005). Additionally, a rough plateau of fMRI and behavioral responses between two duration steps of 120 to 500 ms was reported for stimulus naming (reduced responses at the adjacent lower step of 40 ms; see Fig. 3 of Grill-Spector et al., 2000). While we expect variation by the type of visual process targeted, our results suggest that 80 ms is sufficient for neurotypical adult observers to categorize any natural view of a face in the visual scene (exempting the cases that faces are occluded, artificially degraded, etc.). A practical implication is that studies measuring neural as well as behavioral generic face (vs. object) categorization responses need not present stimuli longer than 80 ms.

Above 80 ms, behavioral accuracy remains at ceiling (above 97% at 6 and 3 Hz; Table 3). Neural amplitude does not increase at these rates: if anything, it decreases (Figure 2A). In the section “Why Are Some Stimuli Not Categorized?”, we discussed how early visual responses may produce masking effects, or competition at a neural level, across neighboring images in RSVP streams. These responses were not specifically represented in the targeted face-categorization responses of the present experiments, which reflect the contrast of faces with non-face objects. However, we also measured general visual responses to stimulus presentation, maximal over the medial occipital cortex, and increasing in amplitude exponentially as stimulus presentation duration increased (Supplementary Figure 1). It is possible these visual responses were more developed as presentation duration increased, being more likely to reach higher-level general visual processing that nevertheless might have interfered more with face-selective processing. Indeed, these stimulus-presentation responses showed some variance in topography across presentation duration, with a particularly distinct response localization at 60 and 120 Hz (Figure 5D), although these latter responses may have been contaminated by noise from the power line frequency at 60 Hz. Thus, one possible explanation for lower amplitudes at longer presentation durations could be an interaction between face and non-face object processing at rates below 12 Hz (compare Supplemental Figure 1 with Figure 2A; and Figure 5B with 5D).

### Individual Differences in Generic Face Categorization

Finding reliable correlations between neural amplitudes and behavioral measures of visual categorization, even when using frequency-tagging with well quantified measures, remains a challenge (e.g., Retter & Rossion, 2016b; Retter & Rossion, 2017; Xu et al., 2017). This is likely because neural responses are heavily influenced by physiological factors: in particular, for EEG the orientation of dipole generators, influenced by individual differences in cortical folding, is thought to play a large role; additional factors include skull thickness and the conductivity of tissue in between the generators and captors (Luck, 2005; Nunez & Srinivasan, 2006; Woodman, 2010).

Here, we developed a method to identify the relationship between individual participants’ behavioral and neural measures. We were able to show a significant correlation between participants’ behavior (measured in Experiment 2 across all presentation rates in terms of inverse efficiency) and face-categorization EEG amplitude (measured in Experiment 1 by weighting the mid-frequency responses from 20–30 Hz (30–50 ms) by that of 3 Hz (333 ms), over the OT ROI; Figure 7D).

This correlation relies on two elements: first, we used a range of presentation rates that accentuated individual differences in generic face categorization. While a great amount of research is being done in terms of individual differences in individual face recognition, differences in generic face vs. object categorization have not been reported to our knowledge, likely due to the high level of expertise that human adults have at performing this function (Crouzet et al., 2010; Herschler & Hochstein, 2005; Scheirer et al., 2014). Here, at presentation rates from 20 to 30 Hz (30 to 50 ms; see Figure 7A and B), we were able to make this function challenging enough to keep all participants below a behavioral ceiling (at 80 ms, i.e., 12 Hz, and later; Figure 2B) and yet above the threshold for measuring significant neural responses in most individual participants (Figure 1).

Second, we used a relative measure of neural activity, weighting individuals’ neural amplitudes in this mid-frequency range by their amplitudes at the lowest presentation rate. At this rate, 3 Hz, individual differences had no relationship to behavioral performance (Figure 7C), again likely due to physiological influences on the neural amplitudes. Using a relative neural measure likely assisted in cancelling out such influences, enabling a more diagnostic signature of individuals’ EEG response amplitudes relative to behavior (again, Figure 7D).

Note again that this relationship is found when comparing explicit behavioral face categorization (Experiment 2, face detection task) to implicit neural face-categorization responses (Experiment 1, naïve, fixation-cross task). Although most (14/16) participants spontaneously reported perceiving faces when describing the sequences after Experiment 1, none reported observing their periodic presentation. Note also that the responses to stimulus-presentation duration showed very different trends, being significant at all durations but decreasing exponentially as duration increased (Supplemental Figure 1), suggesting that the targeted responses are truly specific to face categorization.

### Summary

We have shown that widely variable natural images of faces can be categorized in contrast to non-face objects at presentation rates as short as 17 ms, and optimally at 80 ms. Individual differences were evident within this range, such that individuals’ relative neural amplitudes (across 30-50 ms) were significantly correlated with their behavioral performance (across all presentation rates). While overall the amplitude of face-categorization responses appeared to increase gradually as presentation duration increased, this was completely accounted for by the proportion of discrete “hits” and “misses” in categorization, with categorized faces producing full neural face-categorization responses and uncategorized faces producing no neural face-categorization responses. This provides convergent neural and behavioral evidence that human perceptual (face) categorization is all-or-none.

## Acknowledgments

This research was supported by grants from the Belgian National Foundation for Scientific Research [http://www.fnrs.be/; grant number FC7159 to TR], the European Research Council [https://erc.europa.eu/; grant facessvep number 284025 to BR], the University of Nevada, Reno Integrative Neuroscience Center of Biomedical Research Excellence [https://www.nigms.nih.gov/Research/DRCB/IDeA/pages/COBRE.aspx; grant number P20 GM103650], and the National Eye Institute [https://nei.nih.gov/; grant number EY010834 to MW, and grant number EY023268 to JF]. The funders had no role in study design, data collection and analysis, decision to publish, or preparation of the manuscript. We are grateful to Andrea Conte for creating the stimulation program XPMan, revision 94; to Caroline Michel, for collecting the stimuli; and to Charles Or for generating the color key for plotting electrode location on a scalp map.

