## Supplemental Figure for "All-or-none visual categorization in the human brain"

**Supplemental Figures**

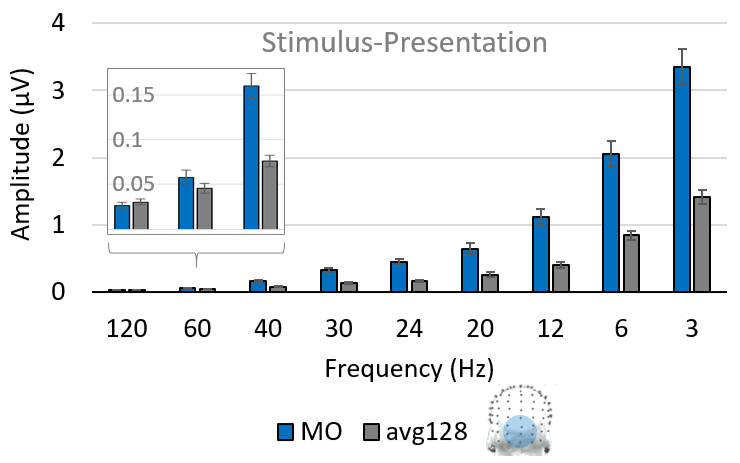
***Supplemental Figure 1****. Quantification of stimulus-presentation responses, correspondingly as in Figure 2B for face-categorization responses****. Key)*** *MO: medial-occipital ROI; avg128 = average of all 128 EEG channels.*

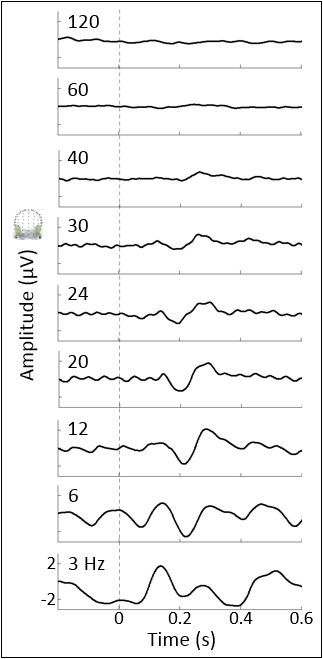
 ***Supplemental Figure 2****. Stimulus-presentation unfiltered time-domain data by condition, with labeling conventions as in Figure 5A. Note that in processing the data were low-pass filtered at 30 Hz, preventing apparent responses at 120-40 Hz, although the amplitude of stimulus-presentation responses was very low at these rates anyway (see Figure 3D and Table 2B).*

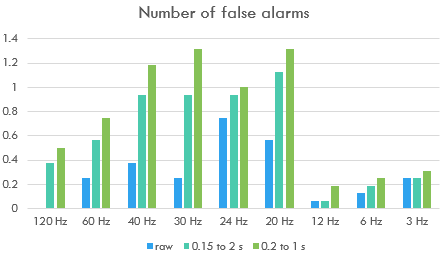

***Supplemental Figure 3****. The mean number of false alarms, summed across all sequence repetitions per frequency (in total about 30 face presentations). Raw = no restrictive time window for correct responses; note that a window of 0.15 to 2 s was used in the analyses, but a more restrictive window of 0.2 to 1 s would have only slightly increased the rate of false positives across frequencies.*
